# Selective Cortical Myelination Reflects Axon-Driven Progression of the BCAS1-positive Oligodendrocyte Lineage

**DOI:** 10.64898/2026.02.15.705957

**Authors:** Huang Zi, Alexander Goldbrunner, Shahrzad Askari, Katharina Eichenseer, Thomas Misgeld, Nicolas Snaidero

## Abstract

Myelination in the cerebral cortex is sparse, discontinuous, showing marked heterogeneity across cortical areas and along individual axons, yet the mechanisms that establish this complex and selective architecture remain unclear. Using histological mapping of oligodendrocyte lineage cells across cortical areas combined with longitudinal in vivo imaging, we find that initial myelination is highly targeted: up to 80 percent of the earliest segments arise consecutively along the same subset of axons. We observed that oligodendrocyte precursor cells enter a premyelinating BCAS1-positive state even within cortical regions that remain poorly myelinated or unmyelinated. However, the progression beyond early premyelinating morphologies correlates with local myelin levels and with interactions with specific neuronal partners. BCAS1-positive ensheathments show a preferential stabilisation and elongation along parvalbumin interneurons compared with somatostatin interneurons, linking regional neuronal composition to differential myelin levels. These findings indicate that cortical myelination emerges from intrinsic lineage programs that are refined through selective interactions with axons, thereby linking neuronal diversity to cortical myelin distribution and providing a framework to understand why remyelination may fail in regions where myelination supportive neuronal populations are reduced.

## Introduction

Myelin in central nervous system is produced by oligodendrocytes and is essential for rapid and precise signal conduction, metabolic support, and stability of cortical circuits^1–4^. In cortex, myelination continues well into adulthood and contributes to experience-dependent plasticity^5–7^. Unlike microglia and astrocytes, which distribute widely and establish near-uniform territorial coverage early in development^8,9^, oligodendrocytes accumulate gradually and follow pronounced spatial and temporal gradients. Primary sensory and motor cortices myelinate early and rapidly, whereas association areas exhibit a more protracted course of integration^10,11^.

In cortex, myelin organization itself is highly complex. Axons differ widely in their myelination patterns, from long continuous stretches to sparse, patchy, or entirely absent internodes^12^. These profiles are not stochastic. During remyelination after oligodendrocyte loss, continuous internodes are rebuilt with single internode precision, whereas patchy profiles are often lost, indicating that cortical myelination is selective and highly constrained^5,13^. How oligodendrocytes select specific axons among thousands of potential targets to generate such axon-specific patterns remains unclear. Oligodendrocytes can initiate wrapping of inert fibers or electrically silent axons once a minimal caliber threshold is exceeded^14,15^, underscoring a strong intrinsic capacity to begin ensheathment independent of axonal cues. Yet axonal diameter alone is not decisive. Thin parvalbumin (PV) interneuron axons, for example, are reliably myelinated ^16^, whereas many larger axons remain unmyelinated^17^, pointing to salient identity-dependent cues for myelination. Several other regulators have been identified that facilitate myelination at different steps of the oligodendrocyte lineage, including activity related signals and axon derived factors that can bias early sheath stabilization^15,18,19^. However, how these influences converge across developmental stages and cortical areas to collectively generate the region- and axon-specific cortical myelin patterns is still unknown.

Oligodendrocyte precursor cells (OPCs), unlike oligodendrocytes, are evenly distributed throughout cortex and therefore cannot account for regional differences in myelin^20,21^. The critical decisions that shape selective myelination are, thus, likely implemented during transitional stages between OPCs and mature oligodendrocytes. However, these stages are rarely captured with the spatial, temporal, and molecular resolution needed, particularly with respect to their interactions with specific axons, to dissect their roles in myelination. Brain enriched myelin associated protein 1 (BCAS1) labels a broad class of premyelinating oligodendrocytes that persist throughout life and display marked morphological diversity, ranging from highly ramified cells to those initiating short sheath-like structures^22,23^. Their position within the lineage, together with the morphological heterogeneity, suggests that BCAS1 positive (BCAS1^+^) cells may represent the stage during which contacts with axons are tested, stabilized, or aborted, thereby shaping the final patterns of cortical myelination, an idea that has not been fully explored^24^.

Cortical myelin is critical in regulating excitation-inhibition balance, temporal precision, and information processing in cortical circuits^25,26^. In the healthy cortex, myelination follows neuron type specific patterns that are tightly linked to functional demands, particularly within fast spiking inhibitory networks including PV interneurons^16,17,27^. These observations highlight that understanding how myelin patterns are established and maintained in the healthy cortex, and how they are selectively organized along certain axons, is directly relevant for elucidating why cortical remyelination would be limited in disease.

Here we combine high resolution histology with longitudinal intravital imaging to characterize how individual oligodendrocytes incorporate into cortical circuits to build myelin specificity at both regional and axonal level. We show that during the initial stage of cortical myelination, oligodendrocytes already exhibit pronounced selectivity, forming the majority of their sheaths consecutively along a restricted subset of axons even when a large pool of unmyelinated targets is present. By charting the oligodendrocyte lineage across cortical regions, we find that BCAS1^+^ cells are broadly generated but diverge markedly in their maturation trajectories between highly and poorly myelinated cortical areas. In sparsely myelinated areas they accumulate in early ramified states, whereas in heavily myelinated regions they progress efficiently toward ensheathing and myelinating states. Finally, by reconstructing interactions between BCAS1^+^ cells and defined neuronal subtypes, we identify a preferential association and elongation of BCAS1^+^ ensheathments along PV interneurons in a BCAS1^+^-cell-stage dependent manner. Together, these findings indicate that cortical myelination patterning is associated with selective incorporation of oligodendrocyte lineage states that is biased by regional context and neuronal identity.

## Results

### Early-onset and persistent regional differences in oligodendrocyte incorporation across cortical areas

To assess myelination dynamics in cortical regions with distinct myelin content, we quantified oligodendrocyte density in *Plp-GFP* mice across cortical areas with anatomically distinct medial-lateral and rostral-dorsal positions as defined by the Allen Brain Atlas (https://atlas.brain-map.org/). These areas included primary motor and sensory cortices (motor cortex (MO), somatosensory cortex (SS), and visual cortex (VIS)), as well as association cortices (temporal association areas (TEa), and lateral entorhinal areas (ENTI) and medial prefrontal cortex (mPFC)) (Fig. 1a). Our analysis focused on the oligodendrocyte cell bodies (Fig. 1b) and spanned a broad timeline, including juvenile (2 weeks; 2W), adolescent (4 weeks; 4W), young adult (2 months; 2M), adult (6 months; 6M), middle-aged (14 months; 14M), and aged (24 months; 24M) mice.

**Figure 1.**
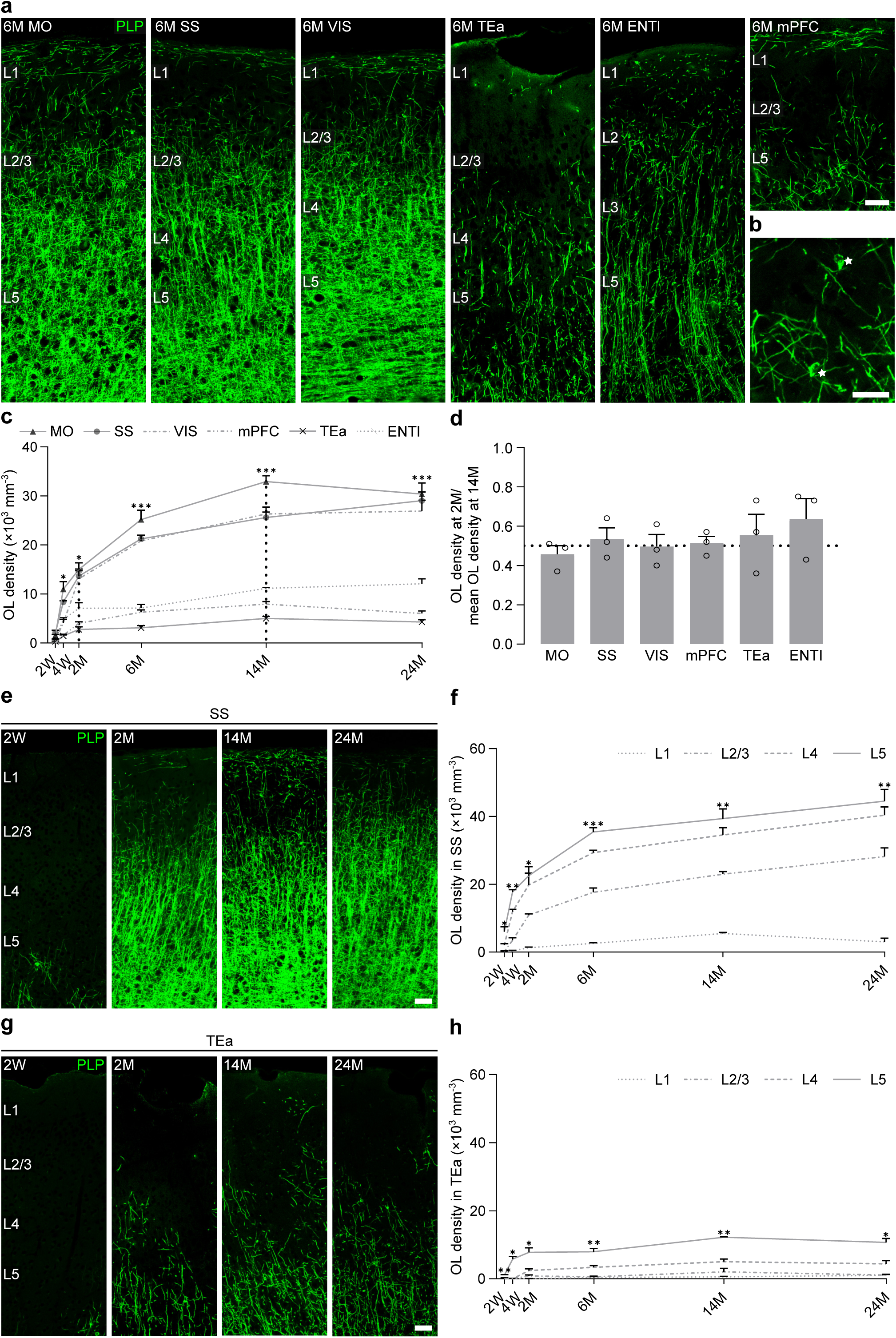
Regional divergences and proportional progression of oligodendrocyte density with development. a, Representative images of 6-month-old *Plp-GFP* mice highlighting regional differences in oligodendrocyte density and myelination in MO, SS, VIS, TEa, ENTl and mPFC. Scale bar, 50 µm; z-projection, 4 µm. b, Representative images showing oligodendrocytes in *Plp-GFP* mice in layer 2/3 of SS. Stars indicate oligodendrocyte cell bodies. Scale bar, 25 µm; z-projection, 4 µm. c, Oligodendrocyte density averaged by cortical layer in each brain area across development (n = 3 animals). Dashed lines indicate the 2M and 14M timepoints. d, Ratio of oligodendrocyte density at 2M relative to the mean density at 14M for each cortical area (n = 3 animals). Dashed line indicates y value = 0.5, corresponding to 50% of the 14M oligodendrocyte density. e, Representative images illustrating myelination in SS at 2W, 2M, 14M and 24M. Scale bar, 50 µm; z-projection, 4 µm. f, Oligodendrocyte density in layer 1, 2/3, 4 and 5 of SS across development (n = 3 animals). g, Representative images illustrating myelination in TEa at 2W, 2M, 14M and 24M. Scale bar, 50 µm; z-projection, 4 µm. h, Oligodendrocyte density in layer 1, 2/3, 4 and 5 of TEa across development (n = 3 animals). Data are presented as mean ± SEM. Repeated-measures (RM) ANOVA is used for comparisons across cortical areas and layers. *P ≤ 0.05, **P ≤ 0.01, ***P ≤ 0.001.

We quantified oligodendrocyte density across cortical layers 1-5 and averaged the data by layer for cross-region comparisons (Fig. 1c). At 2W, oligodendrocyte densities are uniformly low across all areas. During early postnatal development until 2M, primary sensory and motor cortices myelinate rapidly whereas association cortices myelinate at significantly slower rates (Fig. S1a-b). These regional divergences in myelination speed persist, albeit less prominently, beyond 2M (Fig. S1a-b). Hence the early-established regional disparities are largely maintained, resulting in two stable patterns: high oligodendrocyte densities in primary sensory and motor cortices and substantially lower densities in association cortices (Fig. S1c-f). Importantly, by 2M, all cortical areas reach approximately half of their plateau oligodendrocyte density, as observed at 14M (lowest: 0.46 ± 0.04 in MO; highest: 0.64 ± 0.10 in ENTl) (Fig. 1d). This indicate that myelin accumulation progresses proportionally across cortical areas, despite pronounced differences in absolute oligodendrocyte densities.

To further illustrate how these early differences manifest across cortical depth, we selected SS and TEa as representative heavily and sparsely myelinated areas (Fig. 1e-h). These two areas had similar rostral-dorsal position and laminar architecture. Focusing on the detailed myelination levels throughout cortical layers in these areas, oligodendrocyte density exhibits layer-specific differences at all ages in both areas (Fig. 1f, h). When comparing the two brain areas directly, the layer-averaged data show higher oligodendrocyte density in SS from 4W on (Fig. S1g). Consistent with the findings above, at 2M, oligodendrocyte density in most cortical layers (layers 2-5) of both regions reached around half of the peak density observed at 14M (lowest: 0.41 ± 0.16 in layer 2/3 of TEa; highest: 0.63 ± 0.11 in layer 5 of TEa) (Fig. S1h).

Together, these findings demonstrate that cortical myelination follows a biphasic trajectory: an early phase of rapid, regionally divergent oligodendrocyte incorporation that establishes the relative scaling of cortical myelination within the first two postnatal months, followed by a slower phase of proportional growth into adulthood that preserves the area-specific myelination.

### Targeted and consecutive myelin sheath integration from initial stages of cortical myelination

Regional divergences of oligodendrocyte density are accompanied by complex patterns of myelin distribution along axons^12^. To determine whether these patterns reflect non-random myelin placement along axons, or instead arise from progressive random filling of unmyelinated axons, we examined the spatial organization of each sheath from individual oligodendrocytes across regions and ages. We focused on myelin sheath patterns, classifying them as either consecutive (adjacent to at least one other sheath) or isolated (without a neighboring sheath) (Fig. 2a). The presence of consecutive sheaths was used as a readout of non-random myelin distribution^5,13,28^.

**Figure 2.**
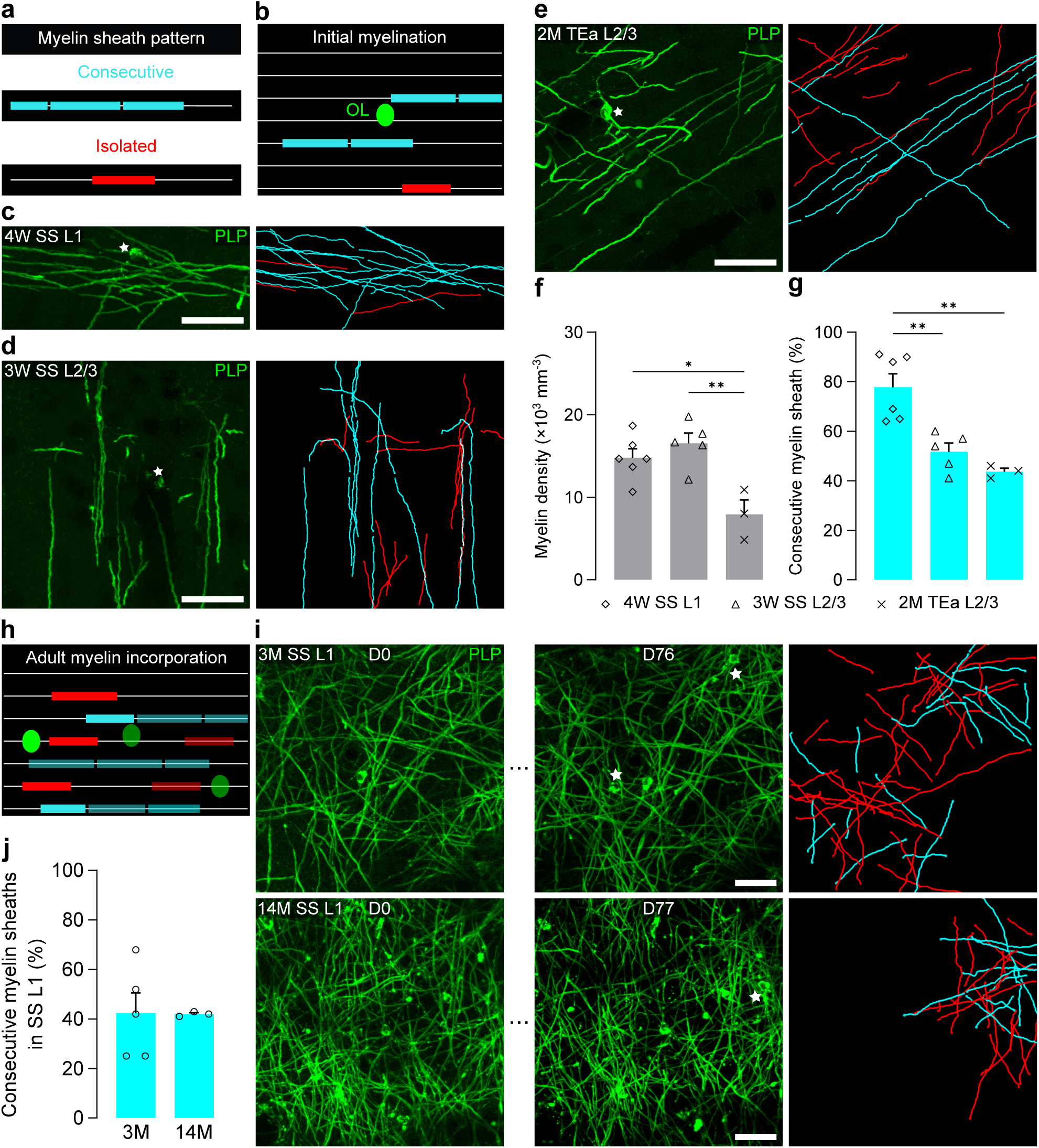
Selective myelin integration across development and adult oligodendrocyte incorporation. a, Schematic illustrating myelin sheath types. Cyan, consecutive sheath (with one or two neighboring sheaths); red, isolated sheath (with no neighboring sheath). Myelin sheaths whose type cannot be defined are excluded from sheath-type classification (unapplicable). b, Schematic illustrating initial myelination. OL, oligodendrocyte in green. c, Left: maximum-intensity projection of initial myelination within a selected 120 × 50 × 40 μm^3^ volume in layer 1 of SS at 4W. Right: projection of 3D reconstruction of myelin sheaths identified in the left panel. Scale bar, 30 µm. d, Left: maximum-intensity projection of initial myelination within a selected 120 × 120 × 20 μm^3^ volume in layer 2/3 of SS at 3W. Right: projection of 3D reconstruction of myelin sheaths identified in the left panel. Scale bar, 30 µm. e, Left: maximum-intensity projection of initial myelination within a selected 120 × 120 × 70 μm^3^ volume in the layer 2/3 of TEa at 2M. Right: projection of 3D reconstruction of myelin sheaths identified in the left panel. Scale bar, 30 µm. (c-e, White stars indicate oligodendrocytes cell bodies.) f, Myelin density during initial myelination in different cortical regions. g, Percentage of consecutive sheaths among all applicable myelin sheaths during initial myelination. (f-g, layer 1 of SS at 4W: n = 6 areas from 3 animals, layer 2/3 of SS at 3W: n = 5 areas from 3 animals, layer 2/3 of TEa at 2M: n = 3 areas from 3 animals, one-way ANOVA with Tukey post hoc test) h, Schematic illustrating newly integrated oligodendrocyte during adulthood. Cell and myelin sheaths with lower intensity indicate pre-existing myelin. i, Maximum-intensity projection of adult myelination in layer 1 of SS at 3M (upper) and 14M (lower). Left: first day of in vivo imaging (180 × 180 × 20 μm^3^). Middle: last day of in vivo imaging (180 × 180 × 20 μm^3^). Right: projection of 3D reconstruction of myelin sheaths produced by the spontaneous newly differentiated oligodendrocytes within the full image (203 × 203 × 80 μm^3^). White stars indicate the newly incorporated oligodendrocytes cell bodies. Scale bar, 30 µm. j, Percentage of consecutive sheaths among all applicable myelin sheaths for newly integrated oligodendrocytes during adulthood (3M: n = 5 cells from 3 animals, 14M: n = 3 cells from 3 animals, student t-test). Data are presented as mean ± SEM. *P ≤ 0.05, **P ≤ 0.01.

Initial myelination, established by the first incorporating oligodendrocytes, occurs in the context of abundant unmyelinated axons. We leveraged our detailed understanding of cortical myelination dynamics in highly and poorly myelinated cortical areas (Fig. 1) and focused on layer 1 and 2/3 of SS and TEa to investigate the initial and early incorporation of myelin sheaths along axons (Fig. 2b). The region of interest was framed according to myelin trajectory, which aligns parallel to the pia in layer 1 (Fig. 2c), but extends in all directions in layer 2/3 (Fig. 2d,e). The earliest appearing oligodendrocytes in layer 1 of SS, at 4W, are accompanied by ∼15 × 10^3^ myelin sheaths per cubic millimeter (Fig. 2f). Notably, 77.83 ± 5.35% of these sheaths are arranged consecutively along the same axons (Fig. 2g). In layer 2/3 of SS, the same myelin density is reached at 3W (Fig. 2f). Here, the percentage of consecutive sheaths is significantly lower (51.80 ± 3.46%) (Fig. 2g), and the average sheath length is shorter (Fig. S2a). The layer-specific myelin pattern did not correlate with overall myelin abundance. Supporting this, layer 2/3 of TEa displays comparable sheath lengths and percentage of consecutive sheaths as SS layer 2/3 at 2M (Fig. 2g, Fig. S2a), despite markedly lower myelin density (Fig. 2f). Together, quantitative analysis revealed repeated sheath placement along restricted subsets of axons across cortical regions during early myelination.

To further investigate whether the initial myelin pattern is a transient feature at early age or conserved principle persisting throughout development, we performed the same analysis in layer 1 and 2/3 of SS at later timepoints (Fig. S2b-e). In layer 1 at 2M, still ∼80% (75.50 ± 3.23%) of the myelin sheaths are consecutive (Fig. S2d). Although sheath density was higher in layer 2/3 than in layer 1 at 2M (Fig. S2c), the myelin sheaths in layer 2/3 were consistently less consecutive and shorter (Fig. S2d-e). The cross-age comparisons revealed that, in both layers of SS, neither the percentage of consecutive sheaths nor the average sheath length changed significantly with continuing sheaths adding (Fig. S2f-k), although the percentage of consecutive sheaths in layer 2/3 increased slowly from 51.80 ± 3.46% at 3W to 60.67 ± 1.86% at 2M (Fig. S2k). These data indicate that the non-random and layer-specific patterns of myelin organization are established early and remain largely stable during postnatal development.

To characterize the precise myelin integration pattern during adulthood, we applied weekly longitudinal in vivo imaging for 80 days to capture spontaneously formed oligodendrocytes (Fig. 2h-j). Based on these high-resolution intravital imaging, we then performed 3D reconstruction of individual sheaths from the newly generated cells. We focused on layer 1 of SS from 3-months-old mice, when adult myelination is still ongoing, and 14-months-old mice when adult myelination saturates (Fig. 1f, Fig. 2i). For individual newly generated oligodendrocytes, 42.40 ± 8.23% of the sheaths are consecutive at 3M (Fig. 2j). Despite a higher cell density and almost absent ongoing myelination at 14M, the percentage of consecutive myelin sheaths of the few new oligodendrocytes we could identify remains stable at 42.00 ± 0.58% (Fig. 2j).

Cortical myelination followed layer-specific patterns across regions. Targeted sheath placement was already evident during initial myelination, adapted between cortical areas, and remained detectable into adulthood. Individual oligodendrocytes frequently formed consecutive sheaths along the same axon.

### c-Fos-based neuronal activity and OPC availability are uncoupled from regional differences in cortical myelination

Neuronal activity is a recognized modulator of myelination, promoting OPC proliferation and differentiation and biasing axonal selection^29,30^. Task-evoked c-Fos has been used to mark neurons that preferentially receive myelin^7^. We therefore ask whether regional or laminar disparities in cortical myelination scale with baseline activity differences, using c-Fos as a general activity readout. Under standard housing with identical handling before perfusion, we quantified NeuN^+^ and c-Fos^+^ neurons in SS and TEa (Fig. 3a-b). Neuronal number is comparable between these two areas (Fig. 3c). We observe a slight, largely non-significant lower number of c-Fos^+^ neurons in TEa, compared with SS throughout the timepoints we analyzed (2W, 4W, 2M, 14M and 24M, Fig. 3d). Furthermore, when calculating the ratio of c-Fos^+^ neurons among the total number of NeuN^+^ neurons, the ratio was broadly similar between these two areas (Fig. S3a). Altogether, the baseline neuronal activity, assessed by c-Fos expression, do not scale with the up to sevenfold difference in myelin content observed at 24M between these areas (SS: 29.03 ± 1.79 × 10^3^ cells mm^−3^; TEa: 4.32 ± 0.44 × 10^3^ cells mm^−3^) (Fig. S1g).

**Figure 3.**
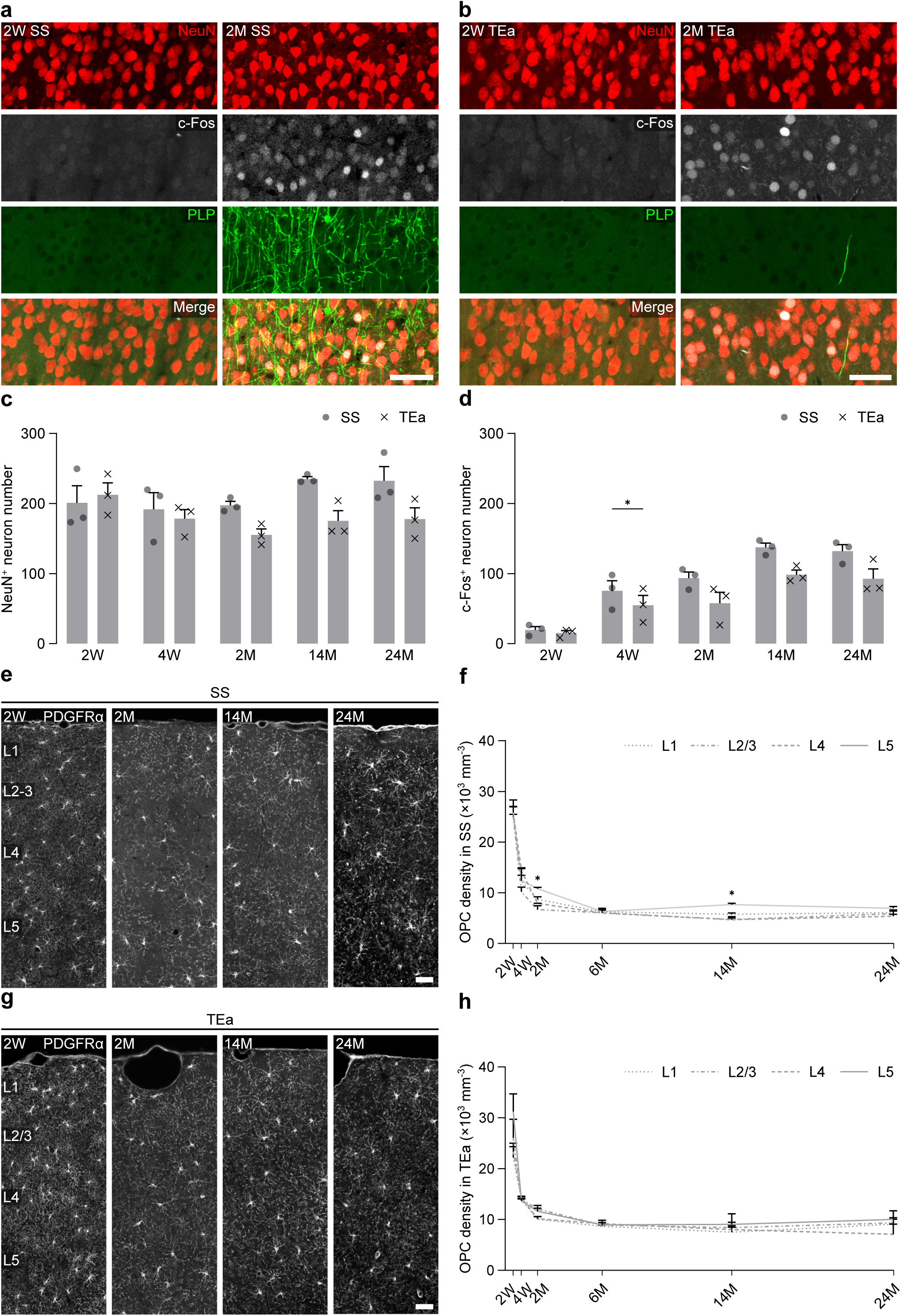
Distribution of c-Fos⁺ neurons and OPCs across cortical regions with distinct myelination levels. a-b, Representative images of NeuN, c-Fos labeling and myelination in *Plp-GFP* mice in SS (a) and TEa (b). Scale bar, 50 µm; z-projection, 4 µm. c, Number of NeuN^+^ neurons within a 250 × 100 × 20 µm^3^ volume in SS and TEa at 2W, 4W, 2M, 14M and 24M (n = 3 animals, multiple paired t-tests). d, Number of c-Fos^+^ neurons within a 250 × 100 × 20 µm^3^ volume in SS and TEa at 2W, 4W, 2M, 14M and 24M (n = 3 animals, multiple paired t-tests). e, Representative images of OPC distribution in SS at 2W, 2M, 14M, 24M. Scale bar, 50 µm; z-projection, 4 µm. f, OPC density in layer 1, 2/3, 4 and 5 of SS across development (n = 3 animals, RM ANOVA). g, Representative images of OPC distribution in TEa at 2W, 2M, 14M, 24M. Scale bar, 50 µm; z-projection, 4 µm. h, OPC density in layer 1, 2/3, 4 and 5 of TEa across development (n = 3 animals, RM ANOVA). Data are presented as mean ± SEM. *P ≤ 0.05.

Having found that c-Fos expression does not mirror the regional distribution of oligodendrocytes, we next examined OPC density. Although OPCs are generally thought to be evenly distributed across the cortex throughout life^21^, we reasoned that a fine-grained analysis across layers and cortical areas might reveal differences relevant to myelination dynamics. We labeled OPCs with PDGFR-α and quantified their density in layers 1-5 of SS and TEa (Fig. 3e-h). OPCs are broadly uniform across layers, except for a transient enrichment in layer 1 of SS at 2M and 14M (Fig. 3f). When data are averaged by layer, TEa exhibits a slightly higher OPC density from 4W onward, reaching significance at 2M (Fig. S3b). Notably, OPC density is either comparable across regions with markedly different myelin levels, or higher in sparsely myelinated areas.

During the period of rapid oligodendrocyte incorporation, spanning from 2W to 2M, OPC density declines sharply across both cortical areas. OPC numbers decrease from ∼25 × 10^3^ cells mm^−3^ at 2W (SS: 25.40 ± 1.52 × 10^3^ cells mm^−3^; TEa: 25.73 ± 2.52 × 10^3^ cells mm^−3^) to ∼10 × 10^3^ cells mm^−3^ by 2M (SS: 8.53 ± 0.14 × 10^3^ cells mm^−3^; TEa: 10.97 ± 0.27 × 10^3^ cells mm^−3^) (Fig. S3b). This pronounced reduction occurs in parallel in regions with high and low myelin content and does not mirror the divergent trajectories of oligodendrocyte accumulation and myelination observed during the same time window (Fig. 1).

Thus, baseline neuronal activity, neuronal density, local OPC availability and the marked postnatal decline in OPC density do not scale with regional differences in cortical oligodendrocyte distribution or myelin abundance.

### The maturation but not generation of premyelinating oligodendrocytes align with regional differences in cortical myelination

Premyelinating oligodendrocytes represent an intermediate stage between OPCs and mature oligodendrocytes^31^. To investigate how these cells distribute relative to regional myelination patterns, we labeled premyelinating oligodendrocytes with BCAS1 antibody and quantified the cell density across cortical layers in SS and TEa throughout development (Fig. 4a,b). In line with previous reports^22^, the density of BCAS1^+^ cells decreases gradually from 2W onward (Fig. S4a,b). In SS, BCAS1^+^ cells are significantly more abundant in the more myelinated deeper layers (layers 4-5) than in superficial layers (layers 1-3) between 2W and 2M (Fig. 4c, Fig. S4c-e). Whereas in TEa this laminar difference is limited to layer 2-4 versus layer 5 at 4W (Fig. S4f,g). When averaged by layer, BCAS1^+^ cell density is higher in SS than in TEa at 4W and 2M, when rapid myelination occurs in SS (Fig. S4h).

**Figure 4.**
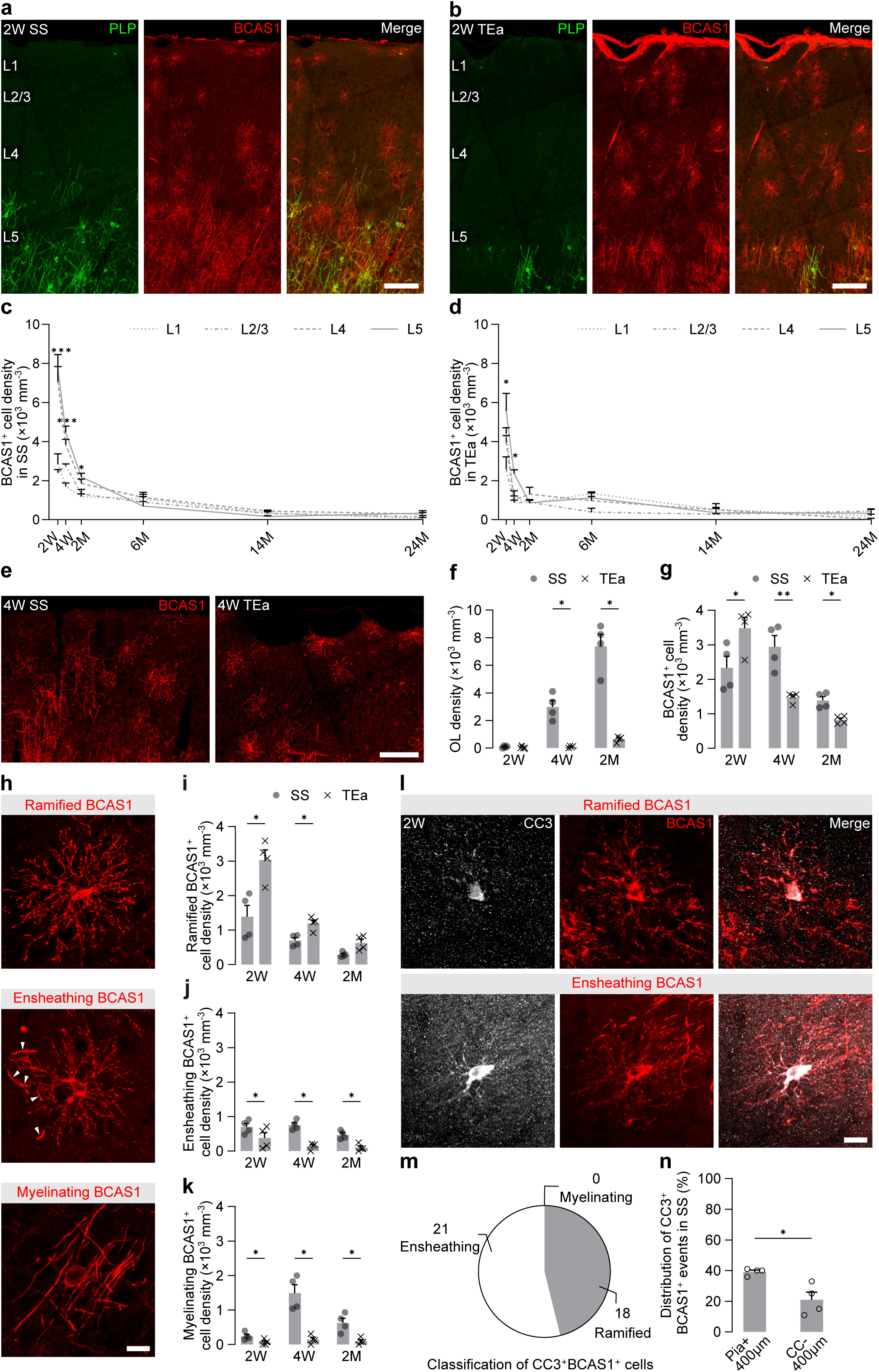
Distinct myelination-associated states of BCAS1^+^ oligodendrocyte lineage cells. a-b, Representative images of myelination and BCAS1 labeling in *Plp-GFP* mice in SS (a) and TEa (b). Scale bar, 100 µm; z-projection, 20 µm. c-d, BCAS1⁺ cell density in layer 1, 2/3, 4 and 5 of SS (c) and TEa (d) across development (2W to 2M: n = 4 animals, 6M to 14M: n = 3 animals, RM ANOVA). e, Representative images of BCAS1 labeling in the upper 400 µm of cortex (from the pia surface) in SS and TEa at 4W. Scale bar, 100 µm; z-projection, 20 µm. f-g, Oligodendrocyte density (f) and BCAS1^+^ cell density (g) in the upper 400 µm cortex of SS and TEa (n = 4 animals, multiple paired t-tests). h, Representative images of ramified, ensheathing and myelinating BCAS1^+^ cells. Scale bar, 15 µm; z-projection, 50 µm. White arrowheads indicating covering of axon by ensheathing BCAS1^+^ cell. i-k, Density of ramified (i), ensheathing (j) and myelinating (k) BCAS1^+^ cells in the upper 400 µm of SS and TEa (n = 4 animals, multiple paired t-tests). l, Representative images of ramified (upper) and ensheathing (lower) CC3^+^BCAS1^+^ cells at 2W. Scale bar, 15 µm; z-projection, 15 µm (upper), 30 µm (lower). m, Classification and quantification of CC3^+^BCAS1^+^ cells exhibiting discernible morphologies. n, Percentage of CC3^+^BCAS1^+^ events located in the upper (400 µm below pia) and lower (400 µm above the corpus callosum) part of SS at 2W (n = 4 animals, paired t-test). Data are presented as mean ± SEM. *P ≤ 0.05, **P ≤ 0.01, ***P ≤ 0.001.

Unlike OPCs, BCAS1^+^ cells are more numerous in regions undergoing active myelination. Nevertheless, the relatively modest regional variations in BCAS1^+^ cell density compared with the pronounced differences in myelination (Fig. S4h, Fig. 1e, g, Fig. S1g) suggest that BCAS1^+^ cell abundance alone does not account for regional myelin levels, prompting further analysis of BCAS1^+^ cell states.

We next focused on the upper 400 µm of cortex in SS and TEa between 2W and 2M, which exhibited marked myelination differences and contained sufficient cells for detailed BCAS1^+^ cell phenotyping (Fig. S4i, Fig. 4e). At 2W, oligodendrocytes are virtually absent in both regions. Thereafter, oligodendrocytes incorporate rapidly in SS, whereas in TEa the incorporation is delayed until 2M, reaching only ∼10% of the oligodendrocyte density in SS (SS: 7.38 ± 0.87 × 10^3^ cells mm^−3^; TEa: 0.65 ± 0.10 × 10^3^ cells mm^−3^) (Fig. 4f). In contrast, differences in BCAS1^+^ cell density are consistently modest, with approximately twofold higher densities in SS than in TEa at 4W and 2M (4W SS: 2.94 ± 0.32 × 10^3^ cells mm^−3^; 4W TEa: 1.25 ± 0.08 × 10^3^ cells mm^−3^; 2M SS: 1.39 ± 0.11 × 10^3^ cells mm^−3^; 2M TEa: 0.83 ± 0.06 × 10^3^ cells mm^−3^), and even higher densities in TEa at 2W (Fig. 4g).

To investigate whether distinct BCAS1^+^ cell states better scale with regional differences in myelination, we then classified BCAS1^+^ cells based on their morphologies: ramified BCAS1^+^ cells, with only extensive processes, representing early differentiation; ensheathing BCAS1^+^ cells, displaying both processes and sheath-like structures; and myelinating oligodendrocytes, displaying only sheath-like extensions, most closely resembling mature oligodendrocytes (Fig. 4h). The density of ramified BCAS1^+^ cells is approximately twofold higher in TEa compared with SS (2W SS: 1.39 ± 0.32 × 10^3^ cells mm^−3^; 2W TEa: 3.03 ± 0.29 × 10^3^ cells mm^−3^; 4W SS: 0.70 ± 0.08 × 10^3^ cells mm^−3^; 4W TEa: 1.20 ± 0.10 × 10^3^ cells mm^−3^; 2M SS: 0.29 ± 0.04 × 10^3^ cells mm^−3^; 2M TEa: 0.63 ± 0.10 × 10^3^ cells mm^−3^), opposite to the regional myelination differences (Fig. 4i). In contrast, we found the densities of ensheathing and myelinating BCAS1^+^ cells are consistently higher in SS at all ages examined, with an up to fivefold difference for ensheathing BCAS1^+^ cells (SS: 0.75 ± 0.07 × 10^3^ cells mm^−3^; TEa: 0.14 ± 0.05 × 10^3^ cells mm^−3^) and tenfold difference for myelinating BCAS1^+^ cells (SS: 1.49 ± 0.25 × 10^3^ cells mm^−3^; TEa: 0.14 ± 0.06 × 10^3^ cells mm^−3^) observed at 4W (Fig. 4j-k).

Previous work has shown that premyelinating oligodendrocytes that fail to integrate are eliminated via apoptosis^32,33^. To assess whether early BCAS1^+^ cell apoptosis relates to subsequent myelination, we co-labeled BCAS1 and cleaved caspase-3 (CC3) at 2W. CC3 immunoreactivity was detected in cells and clustered puncta, and both were included in our analysis. More than half of the apoptotic events are BCAS1^+^ in both SS and TEa (SS: 66.75 ± 3.17%; TEa: 63.25 ± 9.68%) (Fig. S4j). Among the 39 CC3^+^BCAS1^+^ cells with identifiable morphologies we analyzed across cortex (Fig. 4l), approximately half exhibit a ramified morphology and half an ensheathing morphology (ramified: 18 cells; ensheathing: 21 cells), whereas no myelinating BCAS1^+^ cells are observed (Fig. 4m).

We next analyzed the spatial distribution of BCAS1^+^ apoptotic events in SS to further explore whether apoptosis contributes to the regional differences in myelination and found that apoptotic events were less frequent adjacent to the corpus callosum (Fig. S4k). Among all BCAS1^+^ apoptotic events detected in SS, ∼ 40% occur in the superficial cortex (pia to 400 µm depth; 39.25 ± 1.11%), compared with roughly 20% in the deeper cortex (400 µm above the corpus callosum; 21.00 ± 4.97%) (Fig. 4n). In TEa, no clear spatial bias is detected with comparable percentage of apoptotic events in superficial and deeper cortex (pia to 400 µm depth: 41.25 ± 14.85%; 400 µm above the corpus callosum: 46.00 ± 15.42%) (Fig. S4l-m). These observations are consistent with higher incorporation efficiency of BCAS1^+^ cells in regions with substantially greater myelination demand.

Together, these findings reveal marked regional heterogeneity within the BCAS1^+^ populations. While ensheathing and myelinating BCAS1^+^ states scale with local myelination levels, ramified BCAS1^+^ cells accumulate disproportionately in sparsely myelinated regions, indicating that regional differences in cortical myelination emerge at the level of BCAS1^+^ cell maturation rather than initial generation.

### BCAS1⁺ ensheathment stabilization is biased toward parvalbumin interneurons and predicts regional myelin density

The data above suggest that the transition of ramified BCAS1^+^ cells into forming sheath-like structures correlates with local myelination demands. In cortex, neurons are not myelinated uniformly; for instance PV neurons are more myelinated than somatostatin (SST) neurons^17,27^. To investigate how BCAS1^+^ ensheathments interact with these two neuronal subtypes, we generated two triple transgenic mouse lines: *PV-Cre* × *tdTomato* × *Plp-GFP* and *SST-Cre* × *tdTomato* × *Plp-GFP*. We performed 3D reconstruction of 69 BCAS1^+^ cells with sheath-like structures in layer 2/3 of SS at 2M, when PV and SST interneurons have reached maturity^34,35^. We further categorized BCAS1^+^ cells based on the average length and number of BCAS1^+^ ensheathments (Fig. 5a-b). Cells in the *initial* group display less than 30 short ensheathments with average length below 17 µm. Cells in the *expansion* group extend 30 or more ensheathments with an average length below 17 µm. Cells in the *stabilization* group are characterized by significantly longer ensheathments with an average length equal to or greater than 17 µm (Fig. S5a-b). The morphological features of BCAS1^+^ cells are comparable between the two mouse lines (Fig. S5c-d).

**Figure 5.**
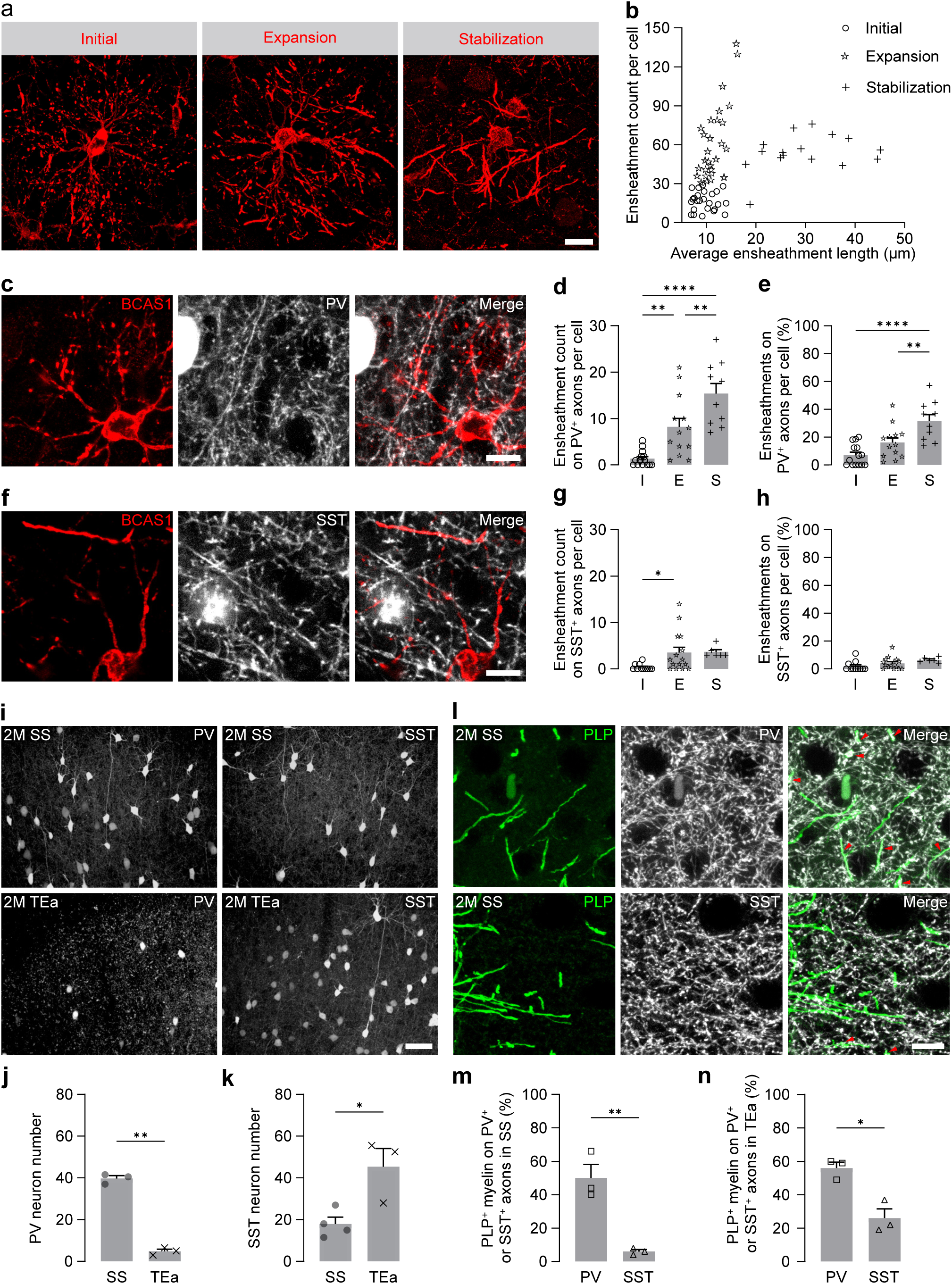
Neuron-specific interactions between axons and BCAS1^+^ ensheathments linked to myelin distribution across brain areas. a, Representative images showing BCAS1^+^ cells from initial, expansion and stabilization groups. Scale bar, 15 µm; z-projection, 20 µm. b, Classification of BCAS1^+^ cells into *initial*, *expansion* and *stabilization* groups based on the number and average length of BCAS1^+^ ensheathments per cell. c, Representative images of BCAS1^+^ ensheathments on PV^+^ axons in *PV-Cre* × *tdTomato* × *Plp-GFP* mice. Scale bar, 10 µm; z-projection, 1 µm. d, Number of BCAS1^+^ ensheathments on PV^+^ axons for individual BCAS1^+^ cells. e, Percentage of BCAS1^+^ ensheathments on PV^+^ axons among all the BCAS1^+^ ensheathments for individual BCAS1⁺ cells. (d-e: *initial*: n = 14 cells from 3 animals, *expansion*: n = 13 cells from 3 animals, *stabilization*: n = 10 cells from 3 animals, one-way ANOVA with Tukey post hoc test). f, Representative images of BCAS1^+^ ensheathments on SST^+^ axons in *SST-Cre* × *tdTomato* × *Plp-GFP* mice. Scale bar, 10 µm; z-projection, 1 µm. g, Number of BCAS1^+^ ensheathments on SST^+^ axons for individual BCAS1^+^ cells. h, Proportion of BCAS1^+^ ensheathments on SST^+^ axons among all the BCAS1^+^ ensheathments for individual BCAS1^+^ cells. (g-h: *initial*: n = 11 cells from 3 animals, *expansion*: n = 15 cells from 3 animals, *stabilization*: n = 6 cells from 3 animals, one-way ANOVA with Tukey post hoc test). i, Representative images of PV (left, *PV-Cre* × *tdTomato* × *Plp-GFP* mice) and SST neuron (right, *SST-Cre* × *tdTomato* × *Plp-GFP* mice) in SS and TEa. Scale bar, 50 µm; z-projection, 50 µm. j-k, Number of PV (j) and SST (k) neuron within a 400 × 300 × 50 µm^3^ volume (PV number in SS and TEa, n = 3 animals, SST number in TEa: n = 3 animals, SST neuron number in SS, n = 4 animals, student-t test). l, Representative images of PLP^+^ myelin sheaths along PV^+^ (upper) and SST^+^ (lower) axons in SS. Scale bar, 10 µm; z-projection, 5 µm. Red arrowheads indicating the myelin sheaths on PV^+^ or SST^+^ axons. m-n, Percentage PLP^+^ myelin sheaths on PV^+^ (m) or SST^+^ (n) axons among all the PLP^+^ myelin sheaths in layer 2/3 of SS and TEa (n = 3 animals, student t-test). Data are presented as mean ± SEM. \**P* ≤ 0.05, \*\**P* ≤ 0.01, \*\*\*\**P* ≤ 0.0001.

Focussing on the ensheathments on PV^+^ or SST^+^ axons (Fig. 5c-h), we found that cells in the *initial* group had on average ∼1 ensheathment on PV^+^ axons (1.35 ± 0.44, Fig. 5d). This number increases to ∼8 in the *expansion* group (8.23 ± 1.83) and nearly doubles again in the *stabilization* group (15.40 ± 2.19) (Fig. 5d). The percentage of ensheathments on PV^+^ axons at a single BCAS1^+^ cell level shows a similar trend, rising from ∼7% (7.07 ± 2.09%) in the *initial* group to more than 30% (31.75 ± 4.50%) in the *stabilization* group (Fig. 5e). In contrast, ensheathments on SST^+^ axons show a more limited increase, reaching ∼4 ensheathments (3.53 ± 1.21) in the *expansion* group and remaining stable in the *stabilization* group (3.67 ± 0.49), with the percentage of ensheathments remaining below 7% in all categories (*initial*: 1.82 ± 1.08; *expansion*: 4.06 ± 1.10; *stabilization*: 6.42 ± 0.73) (Fig. 5g-h). These observations indicate that BCAS1^+^ ensheathment patterns differ between axonal subtypes across maturation states.

When comparing the ensheathments on PV^+^ and SST^+^ axons, the number and percentage of ensheathments on PV^+^ axons in both *expansion* and *stabilization* groups are higher than those on SST^+^ axons, although statistical significance is not reached for ensheathment number in the *expansion* group (Fig. S5e-f). In the *expansion* group, approximately twice as many ensheathments are found on PV^+^ axons compared to SST^+^ axons (PV^+^ axons: 8.23 ± 1.83; SST^+^ axons: 3.53 ± 1.21) (Fig. S5e). Together with our finding that PV neurons were approximately twice as numerous as SST neurons within layer 2/3 of SS (PV neurons: 39.67 ± 1.36; SST neurons: 17.88 ± 3.32), these data suggest that contacts of ensheathments at the expansion stage may depend on local axonal availability (Fig. S5g).

Alongside these findings, we observe a substantial number of BCAS1^+^ ensheathments on pre-existing myelin sheaths (Fig. S5h). To examine this further, we analyzed ensheathments formed on PLP^+^BCAS1^−^ mature myelin sheaths in *PV-Cre × tdTomato × Plp-GFP* mice across the three BCAS1^+^ cell groups (Fig. S5i-j) and compared these with ensheathments on PV^+^ axons (Fig. S5k-l). In both *initial* and *expansion* groups, ∼20% of ensheathments are located on existing mature myelin sheaths (initial: 25.69 ± 5.89%; expansion: 21.96 ± 4.31%), exceeding those on PV^+^ axons in the *initial* group (7.07 ± 2.09%) (Fig. S5l). In these two groups, ensheathment number on myelin sheaths positively correlates with local oligodendrocyte density (Fig. S5m), consistent with our idea that local target availability influences ensheathments at early stage. Notably, ensheathments on mature myelin sheaths are largely absent in *stabilization* group cells, with only 1 out of 10 cells displaying such contacts (2 out of 519 ensheathments in total), whereas ensheathments on PV^+^ axons increase in both number and length, highlighting a stage-specific refinement of myelin targeting (Fig. S5k).

Because the myelination along PV^+^ and SST^+^ axons has been reported to occur preferentially near the soma^36^, we quantified the numbers of PV and SST interneurons in layer 2/3 of SS and TEa to assess whether regional differences in neuronal composition correlate with cortical myelin levels (Fig. 5i). We find that PV interneuron number is nearly eightfold higher in SS compared to TEa at 2M (SS: 39.67 ± 1.36; TEa: 4.83 ± 1.01) (Fig. 5j), while SST number shows the opposite trend with approximately threefold more SST neurons in TEa (SS: 17.88 ± 3.32; TEa: 45.33 ± 8.71) (Fig. 5k). These differences persist until 14M, consistent with the regional myelination differences (Fig. S5n-o, Fig. 1). We then examined the distribution of mature myelin along PV and SST neurons in 2-month-old SS and TEa (Fig. 5l). Despite neuronal densities exhibiting opposite regional trends, the proportion of myelin sheaths on PV^+^ axons remained significantly higher than along SST^+^ axons with around tenfold enrichment in SS (PV^+^ axons: 50.24 ± 8.04%; SST^+^ axons: 5.72 ± 1.07%) and approximately twofold in TEa (PV^+^ axons: 55.77 ± 3.37%; SST^+^ axons: 26.16 ± 5.70%) (Fig. 5m-n).

Together, these observations show that preferential stabilization and elongation of BCAS1^+^ ensheathments along PV^+^ axons coincide with regional differences in interneuron composition and cortical myelin levels.

## Discussion

In this study, we combine longitudinal imaging, spatial analyses, and cell state characterization to dissect how early oligodendrocyte lineage dynamics contribute to axon- and region-specific cortical myelin patterning. Our data show that cortical myelination is not a linear consequence of oligodendrocyte differentiation, but it emerges through a staged process involving extensive exploration, selective stabilization, and constrained progression of individual cells, with early bias toward restricted axonal subsets. By quantifying oligodendrocyte incorporation and myelin organization across cortical areas and developmental stages, we show that regional differences in cortical myelination emerge early and are maintained proportionally across adulthood. Across cortical areas that ultimately develop distinct myelin densities, oligodendrocyte lineage cells broadly engage in a premyelinating state, yet only a subset successfully transitions into stable, myelinating oligodendrocytes. At the level of individual cells, this transition is accompanied by highly targeted myelin placement, with repeated formation of consecutive sheaths along restricted axonal subsets from the onset of myelination. This selective and constrained progression is consistent with in vivo work demonstrating repeated, transient ensheathment attempts and a restricted temporal window for successful sheath stabilization^37,38^. Once formed, however, myelin sheaths and the oligodendrocytes that generate them exhibit remarkable long-term stability, underscoring that early selection events exert durable effects on circuit organization^3,39,40^.

A key insight emerging from our analyses is that BCAS1^+^ oligodendrocytes represent as a functional checkpoint within this process. OPCs across cortical regions retain the intrinsic competence to enter the BCAS1^+^ state, including in aging cortex and disease states ^22,41^. While BCAS1^+^ cells have been described as marking regions of active myelin formation^22^, our data show that entry into this state alone does not predict successful myelination. Indeed, sparsely myelinated TEa displays numbers of ramified BCAS1^+^ cells higher than those observed in more heavily myelinated SS, yet these cells show delayed and limited progression toward ensheathing and myelinating states (Fig. 4). This suggests that progression into the BCAS1^+^ premyelinating state is broadly permissive and may reflect intrinsic differentiation programs and cortex-wide cues. Consistent with this interpretation, recent work has shown that OPCs throughout the adult central nervous system repeatedly initiate differentiation with spatial and temporal regularity, largely independent of local myelin demand^42^. Following entry into the BCAS1^+^ state, cells adopt a branched morphology consistent with active environmental sampling^43^, and the emergence of ensheathing or stabilizing states reflects successful engagement with extrinsic, permissive axonal surfaces, scaling with regional myelination levels (Fig. 4). BCAS1 expression should therefore not be interpreted solely as a marker of imminent myelination, but as a vulnerable intermediate state in which cells must successfully resolve local selection pressures to progress, consistent with earlier evidence that axon-derived signals are required for oligodendrocyte survival during differentiation^44^.

Our data further indicate that progression beyond the BCAS1 stage depends on the stabilization of early axonal contacts. Premyelinating oligodendrocytes adopt a branched, exploratory morphology that supports motility and repeated axonal sampling^37^. At intermediate stages, BCAS1^+^ cells form short, dynamic ensheathments, including contacts with mature myelin sheaths and axons of differing myelination propensity, consistent with an exploratory rather than committed state. Indeed, the frequency of these early ensheathments scales with the local abundance of PV and SST neurons and extends to contacts with pre-existing myelin sheaths, which are also engaged by BCAS1^+^ cells at this stage (Fig. 5). Initiation of axonal ensheathments represents a partial commitment that progressively anchors the cell, constraining further relocation and reducing its accessible search space^38^. Stabilization into a mature, myelinating oligodendrocyte is associated with the formation of multiple consecutive, spatially aligned ensheathments. This concept is directly supported by our observation that myelin is placed in a highly targeted manner from the onset of myelination, with up to ∼80% of early sheaths arranged consecutively along a small subset of axons, despite the presence of abundant unmyelinated targets^45^. Importantly, the proportion of consecutive sheaths rarely falls below ∼40% across cortical layers, regions, or ages, including in layer 2/3 of both SS and TEa and in older animals (Fig. 2), indicating that consecutive sheath formation represents a quantitative correlate for stable oligodendrocyte integration rather than a transient developmental feature. While PV^+^ axons are known to be preferentially myelinated^17,36,46,47^, our data indicate that regional PV neuron density critically shapes the probability that individual BCAS1^+^ cells encounter sufficient compatible axonal targets to accumulate the minimum fraction of consecutive ensheathments required for stabilization. Consistent with this, we find that BCAS1^+^ ensheathments preferentially stabilize and elongate along PV axons as cells progress toward more mature states (Fig. 5). Conversely, in sparsely myelinated regions, cells are more likely to enter commitment states without encountering sufficient stabilizing targets, leading to attrition.

In this framework, mentioned stabilization requirements impose state-dependent constraints on cellular search strategy and survival. OPCs are motile cells capable of surveying large tissue volumes, enabling constant sampling of changing potential axonal targets^43,48^. While branched BCAS1^+^ cells may retain some capacity for continued exploration, progression into ensheathment states restricts movement and creates a finite window for successful stabilization^37,38^. Cells that fail to reach threshold within this window are selectively lost^5,23^, consistent with evidence for turnover of differentiating oligodendrocytes and efficient clearance of individual dying cells in vivo^39,49^. In line with this interpretation, we detect apoptotic events predominantly within ramified and early ensheathing BCAS1^+^ states, whereas myelinating BCAS1^+^ cells are largely spared, indicating that failure to stabilize is resolved through selective cell loss rather than prolonged persistence (Fig. 4). Within SS, we observe fewer CC3^+^ cells in deeper layers, where myelination is more extensive, compared with superficial layers (Fig. 4), further supporting a link between stabilization success and local myelination context.

These constraints are expected to become particularly consequential under conditions that increase stabilization demands or reduce access to permissive axonal targets. Neuroinflammatory conditions may not prevent entry into premyelinating programs, as premyelinating oligodendrocytes and BCAS1^+^ cells are present in inflammatory lesions, but instead elevate the requirements for successful stabilization, thereby biasing repair toward selective loss rather than productive remyelination^41,50–53^. In cortical demyelinating disease, neuronal loss, altered activity patterns, and inflammatory signaling converge to reduce the density and quality of myelination-competent axonal targets, particularly within inhibitory networks that are selectively vulnerable^54,55^. In the context of our findings, such changes are predicted to impair progression beyond the BCAS1^+^ ensheathment stage by limiting access to stabilizing axonal partners. Under these conditions, oligodendrocyte lineage cells may still enter premyelinating states but are less likely to stabilize before mobility is lost, consistent with our observation that entry into BCAS1^+^ states is broadly preserved, whereas stabilization and survival depend on successful axonal engagement. This framework provides a mechanistic explanation for why therapeutic strategies that primarily promote OPC differentiation have shown limited efficacy^56^, as such approaches do not address constraints imposed by target availability, spatial organization, or survival during early commitment stages^57–60^. By contrast, our data suggest that effective remyelination strategies should move beyond forcing lineage progression and instead build on the bias of cortical myelination toward specific axonal populations that support efficient search and stabilization. We identify PV interneuron axons as a prominent example of preferential substrates for BCAS1^+^ ensheathment stabilization, highlighting that axonal identity and local target availability are central determinants of successful myelination. In this framework, the primary limitation lies not in lineage entry, but in whether BCAS1^+^ cells encounter sufficient permissive axonal surface within a constrained temporal window. Approaches that increase access to myelination-competent axonal targets, for example through activity-dependent mechanisms or by facilitating effective cell–axon encounters, would be expected to enhance stabilization at this intermediate stage. Beyond PV axons per se, the preferential stabilization and elongation of BCAS1^+^ ensheathments on defined axonal subsets suggest that specific axonal surface molecules or membrane properties confer enhanced permissiveness for oligodendrocyte stabilization, reinforcing that neuronal identity and membrane properties shape stabilization competence.

Together, our findings argue that cortical myelin patterns arise from dynamic, state-dependent selection processes rather than uniform differentiation efficiency. By linking regional myelin outcomes to a defined stabilization checkpoint within the oligodendrocyte lineage, this work provides a framework for understanding regional myelin heterogeneity in health and disease and for guiding interventions that align with the biological constraints of cortical myelination.

## Acknowledgements

This project was supported by funding to N.S. from the Hertie Network of Excellence in Clinical Neuroscience (P1200019) and the Else Kröner-Fresenius-Stiftung (2022_EKEA.162). Additional support to N.S. was provided by the Hertie Foundation as part of the independent research group leader funding scheme. High-resolution microscopy was supported via a DFG instrumentation grant (INST 37-1170-1 FUGG, ID 467868227 and INST95/1755-1 FUGG – ID 518284373). Related work in the laboratory of T.M. is supported by the Deutsche Forschungsgemeinschaft (TRR 274/1-2 2020, project B03 and Z01, ID 408885537, Mi 694/9-1/2 A03, ID 405358801, FOR Immunostroke, TRR 167/3 2025, ID 259373024, and CRC 1744/1 2026, ID 548585053). T.M. also receives support from the Munich Center for Systems Neurology (SyNergy EXC 2145, ID 390857198) and the German Center for Neurodegenerative Diseases. We are grateful to J. Saile, D. Steinmetz, and N. Budak for excellent animal husbandry. We also thank B. Zalc (ICM Paris) for his generous provision of Plp-GFP(z) mice^13^.

## Author contributions

N.S. and Z.H. conceptualized the project and experiments and N.S. provided overall scientific supervision. Z.H. performed most of the investigations, including structural imaging, data curation, and analysed most of the data. A.G., S.A., and K.E. contributed to generating samples, performing experiments and analysing data. N.S. and T.M. contributed to funding acquisition, infrastructure, and T.M. provided scientific input through discussion; N.S. and Z.H. wrote the manuscript with input of all authors.

## Methods

### Animal care and use

Female and male *Plp-GFP* mice^13^ of six ages (2W, 4W, 2M, 6M, 14M and 24M) were used to analyze oligodendrocyte lineage dynamics, neuronal activity and myelin patterns. *Plp-GFP × Cspg4-dsRed* (JAX, 008241) mice at 3M and 14M (both sexes) were used for in vivo imaging related experiments. *PV-Cre* (JAX, 017320) × *tdTomato* (JAX, 007914) *× Plp-GFP* and *SST-Cre* (JAX, 013044) × *tdTomato × Plp-GFP* mice (2M, both sexes) were used for neuron-specific analysis. All mice were healthy and did not display any abnormal behavior phenotypes and were housed in groups < 5 under a 12-h light/dark cycle with ad libitum access to food and water.

All animal experiments were performed in accordance with the regulations of the relevant animal welfare acts (TierSchG) and protocols approved by the respective regional authorities (Regierung von Oberbayern and Baden-Württemberg).

### Immunohistochemistry

Mice were deeply anesthetized with isoflurane and transcranial perfused with 4% paraformaldehyde (PFA; in 0.1 M phosphate buffer, pH 7.4). Brains were post-fixed for 8 h in PFA, transferred to PBS, and cut into coronal sections. Free-floating sections were permeabilized in 1.0% Triton X-100/ PBS (30 min, room temperature) and incubated in blocking solution (0.5% Triton X-100, 10% BSA in PBS, 1 h). For mouse-derived primary antibodies, additional Fc-fragment blocking was performed (1:100, 1 h). Sections were incubated overnight at 4 °C in primary antibody, followed by secondary antibody incubation. Sections were mounted with antifading medium and then imaged. Mouse and rabbit anti-BCAS1 antibodies were used interchangeably depending on host species compatibility in multiplex stainings and showed comparable labeling patterns.

The primary antibodies used in this study were listed below.

| Primary antibody | Host species | Source | Catalog | Dilution |
| --- | --- | --- | --- | --- |
| c-Fos | mouse | Santa Cruz | sc-166940 | 1:300 |
| NeuN | rabbit | Cell signaling technology | #24307 | 1:500 |
| PDGFR-α | rabbit | Cell signaling technology | #3164 | 1:250 |
| BCAS1 | mouse | Santa Cruz | sc-6954 | 1:250 |
| Cleaved caspase 3 | rabbit | Cell signaling technology | #9661 | 1:300 |
| BCAS1 | rabbit | Synaptic system | 445008 | 1:2000 |

### Imaging acquisition and processing

The images were acquired using Olympus FV3000 confocal laser-scanning microscope. For cell density analysis, z-stacks (1024 × 1024 pixels; 2.0 µm step size) were imaged with a ×20 air objective (NA 0.75) and a 2× zoom. For myelin tracing, z-stacks (1024 × 1024 pixels; 0.5 µm step size) were taken with a ×40 oil objective (NA 0.95) and 1.7× zoom. For BCAS1^+^ ensheathments analysis, z-stacks (1024 × 1024 pixels; 0.5 µm step size) were taken with a ×63 oil objective (NA 0.95) and 1.7× zoom. Analysis was then performed in Fiji with cell counter or SNT plugin^61,62^, depending on the dataset. Image analysis and quantification were performed blinded to brain area, age, and genotype.

Chronic in vivo imaging of the layer 1 oligodendrocytes and myelin segments was initiated 14-21 days after the implantation of the cranial window and was performed using an Olympus FV-RS microscope (Olympus, Japan) equipped with femtosecond pulsed Ti:Sapphire lasers (Insight and Mai Tai HP-DS, Spectra-Physics) at a maximum power of 30 mW (measured the back focal plane). Imaging was performed with a resonant scanner using 16× averaging.

### Myelin tracing

The images were processed with Fiji. ROIs for analysis were defined according to the orientation of myelin sheaths. For layer 1, areas of 50 × 150 × 40 µm^3^ with long axis in parallel to pia was selected. For layer 2/3 of somatosensory cortex, an area of 120 × 120 × 20 µm^3^ was framed to have comparable volume for analysis as layer 1. For layer 2/3 of temporal association areas, we increased the size to 120 × 120 × 70 µm^3^ to have enough number of myelin sheaths for analysis. All sheaths crossing the ROI were traced in Fiji using simple neurite tracer until their ends were defined or terminated within the imaged areas. The myelin sheaths were categorized as isolated and consecutive. Isolated sheaths did not have neighbouring sheaths within 5 µm in both ends. Consecutive sheaths had at least one neighbour myelin sheaths, including sheaths with one end adjacent and other extending beyond the imaged areas. The 5 µm threshold was chosen to exceed the axial resolution of the imaging system and to reflect internodal spacing beyond maximum node of Ranvier length. Sheaths with both ends extended beyond the imaging boundaries were classified as ‘unapplicable’ and these were excluded from the sheath type analysis.

### Cranial window implantation

To enable longitudinal imaging, a 4-mm cranial window was implanted above the somatosensory cortex as previously described^13^. In short, mice were anaesthetized with a mixture of medetomidine (0.5 mg/kg), midazolam (5 mg/kg) and fentanyl (0.05 mg/kg) intraperitoneally. A craniotomy was performed with a 0.5-mm drill head (Meisinger), and the window was secured with dental cement.

### In vivo imaging and myelin tracing

On the imaging days, animals were anesthetized with 2% isoflurane for induction and maintained at 1.2–1.5% for the rest of the imaging period. Physiological parameters (anesthesia depth, blood oxygenation, heart rate) were continuously monitored with a MouseOx system (Starr Life Science Corp). On the first imaging day, several areas (5–8; *x*/*y*/*z*: 203/203/90 µm^3^) were randomly selected and imaged with a pixel size of 0.4 µm in *x*/*y* and 0.5 µm in *z*. The same areas were imaged every 3–7 days for up to 80 days. We observed no signs of photo-damage over the imaging period using these imaging conditions. Areas showing deteriorated imaging quality or without new oligodendrocyte incorporation were excluded.

Analysis was performed on PLP channel only. Only myelin sheaths located along the trajectory of newly formed oligodendrocytes and appearing within 2 weeks after the initial detection of the oligodendrocyte cell body (PLP expression) were included in the analysis. The myelin sheaths were traced in Fiji using *Simple Neurite Tracer* and classified as isolated or consecutive as described above.

### Statistics

For all analyses, individual animals were treated as the biological replicate. When multiple imaging areas or cells were analyzed from the same animal, values were averaged per animal prior to statistical comparison unless stated otherwise. Statistical analyses were performed with Prism 10.0. Each figure legend contained sample size, statistical tests, and the corresponding significance level (p value). Data were reported as mean ± SEM. Statistical significance is indicated as follows: *P* ≤ 0.05 (*)*, P ≤ 0.01* (**)*, P ≤ 0.001* (***), and *P* ≤ 0.0001 (****). Data distribution was assessed prior to the application of parametric statistical tests.

### Use of generative AI

We used large-language model (LLM)-based software (ChatGPT 5) as language editing tool. All results were checked and edited by the authors. No data analysis or figure material was generated using LLMs.

### Data availability

The datasets generated and analyzed during the current study are available from the corresponding authors on reasonable requests. Mouse lines can be requested under academic material transfer agreements from The Jackson Laboratories or from the laboratories that generated the lines, as indicated.

---

**Supplementary figure 1.**
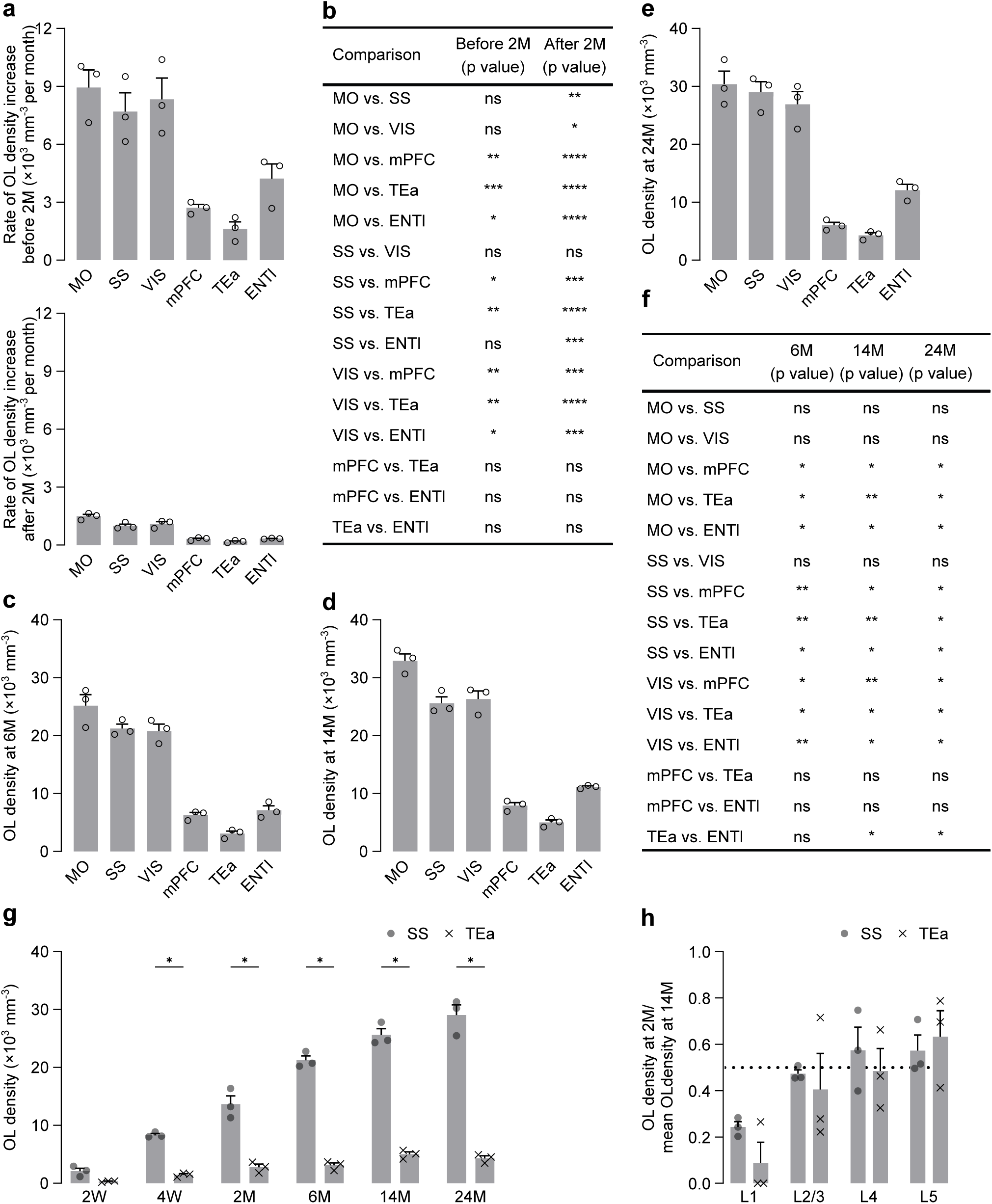
Region- and layer-specific developmental dynamics of oligodendrocyte density in the cortex. a, Rate of increase in oligodendrocyte density per month in different brain areas before (upper) and after (lower) 2M. b, Comparisons of oligodendrocyte density increasing rate between different brain areas before and after 2M (n = 3 animals, RM ANOVA with Tukey post hoc test). c-e, Oligodendrocyte density averaged by cortical layer in different brain areas at 6M (c), 14M (d) and 24M (e). f, Comparisons of layer-averaged oligodendrocyte density between different brain areas at 6M, 14M and 24M (n = 3 animals, RM ANOVA with Tukey post hoc test). g, Oligodendrocyte density averaged by cortical layer in SS and TEa across development (n = 3 animals, multiple paired t-tests). h, Ratio of oligodendrocyte density at 2M relative to the mean density at 14M for layer 1, 2/3, 4 and 5 in SS and TEa (n = 3 animals, multiple paired t-tests). Dashed line indicates y value = 0.5, corresponding to 50% of the 14M oligodendrocyte density. Data are presented as mean ± SEM. *P ≤ 0.05, **P ≤ 0.01, ***P ≤ 0.001, ****P ≤ 0.0001.

**Supplementary figure 2.**
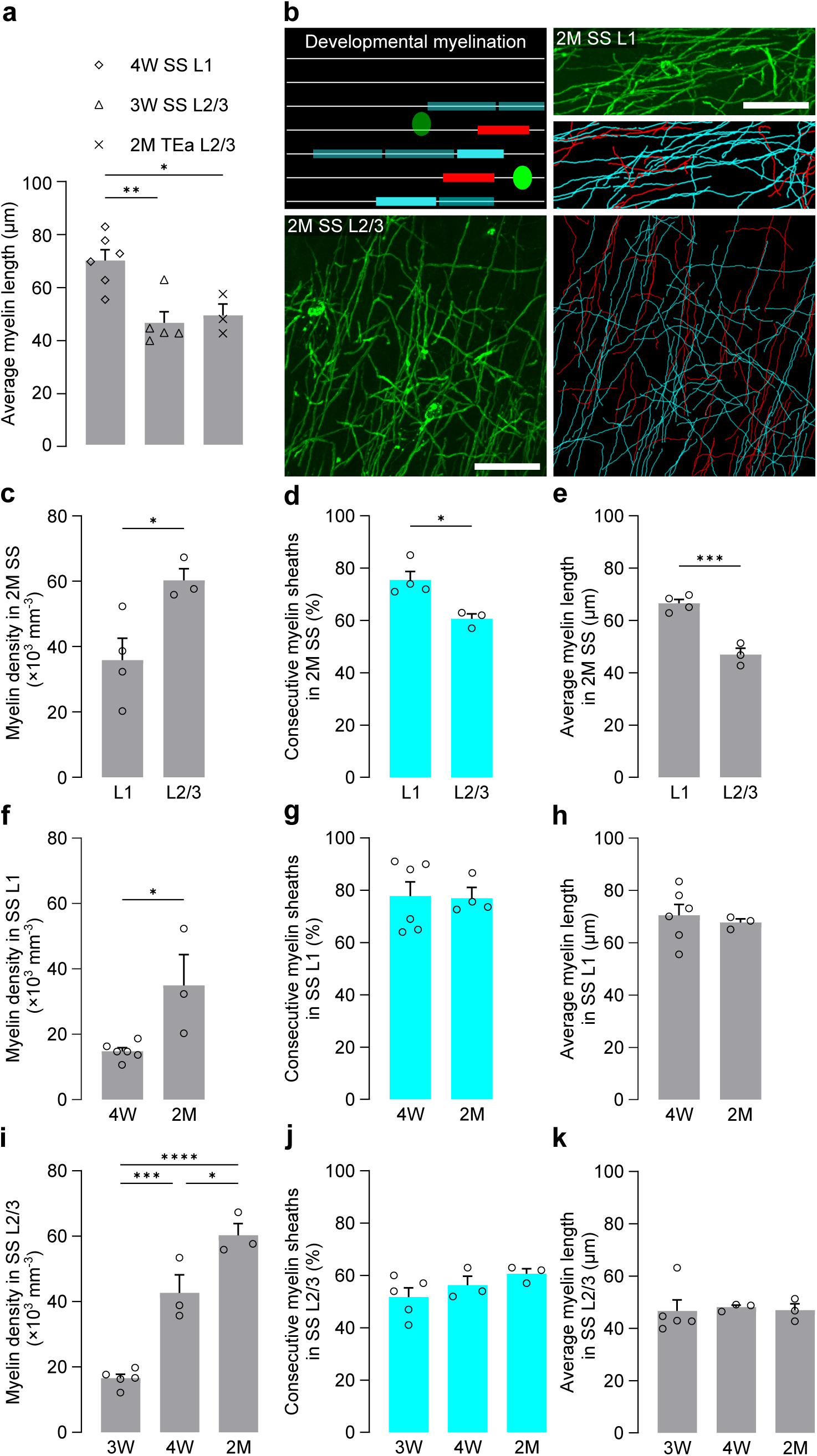
Persistence of early-established myelin patterns with development. a, Average sheath length for initial myelination in different cortical regions (layer 1 of SS at 4W: n = 6 areas from 3 animals, layer 2/3 of SS at 3W: n = 5 areas from 3 animals, layer 2/3 of TEa at 2M, n = 3 areas from 3 animals, one-way ANOVA with Tukey post hoc test). b, Schematic, maximum-intensity projection and 3D reconstruction of developmental myelination in layer 1 (120 × 50 × 40 µm^3^) and layer 2/3 (120 × 120 × 20 µm^3^) of SS at 2M. Cell and myelin sheaths with lower intensity indicate pre-existing myelin. Scale bar, 30 µm. c, Myelin density in layer 1 and layer 2/3 of SS at 2M. d, Percentage of consecutive myelin sheaths among all the applicable sheaths in layer 1 and layer 2/3 of SS at 2M. e, Average sheath length in layer 1 and layer 2/3 of SS at 2M. (c-e, layer 1 of SS at 2M: n = 4 areas from 3 animals, layer 2/3 of SS at 2M: n = 3 areas from 3 animals, Student t-test). f, Myelin density in layer 1 of SS at 4W and 2M. g, Percentage of consecutive myelin sheaths among all the applicable sheaths in layer 1 of SS at 4W and 2M. h, Average sheath length in layer 1 of SS at 4W and 2M. (f-h, layer 1 of SS at 4W: n = 6 areas from 3 animals, layer 1 of SS at 2M: n = 3 areas from 3 animals, Student t-test). i, Myelin density in layer 2/3 of SS at 3W, 4W and 2M. j, Percentage of consecutive myelin sheaths among all the applicable sheaths in layer 2/3 of SS at 3W, 4W and 2M. k, Average sheath length in layer 2/3 of SS at 3W, 4W and 2M. (i-k, layer 2/3 of SS at 3W: n = 5 areas from 3 animals, layer 2/3 of SS at 4W: n = 3 areas from 3 animals, layer 2/3 at 2M: n = 3 areas from 3 animals, one-way ANOVA with Tukey post hoc test). Data are presented as mean ± SEM. *P ≤ 0.05, **P ≤ 0.01, ***P ≤ 0.001, ****P ≤ 0.0001.

**Supplementary figure 3.**
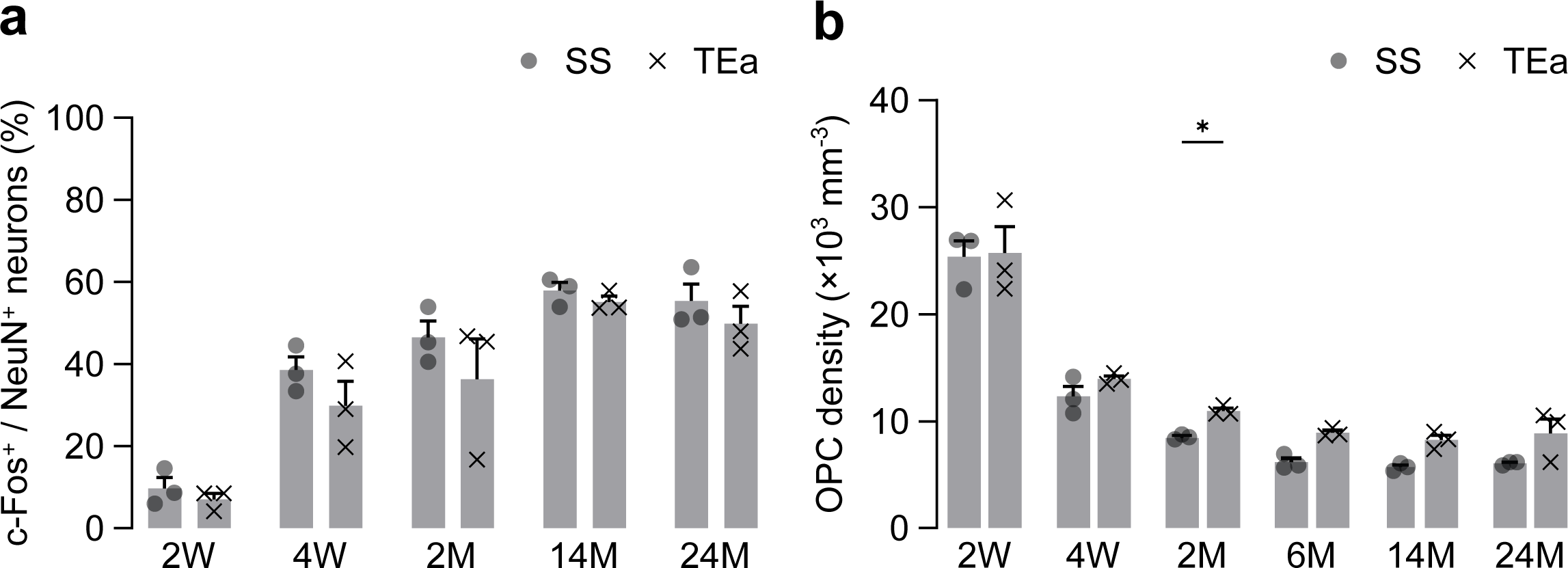
Fraction of c-Fos^+^ neurons and OPC density in SS and Tea. a, Percentage of c-Fos^+^ neurons among NeuN^+^ neurons in SS and TEa at 2W, 4W, 2M, 14M and 24M (n = 3 animals, multiple paired t-tests). b, OPC density averaged by cortical layer in SS and TEa across development (n = 3 animals, multiple paired t-tests). Data are presented as mean ± SEM. *P ≤ 0.05.

**Supplementary figure 4.**
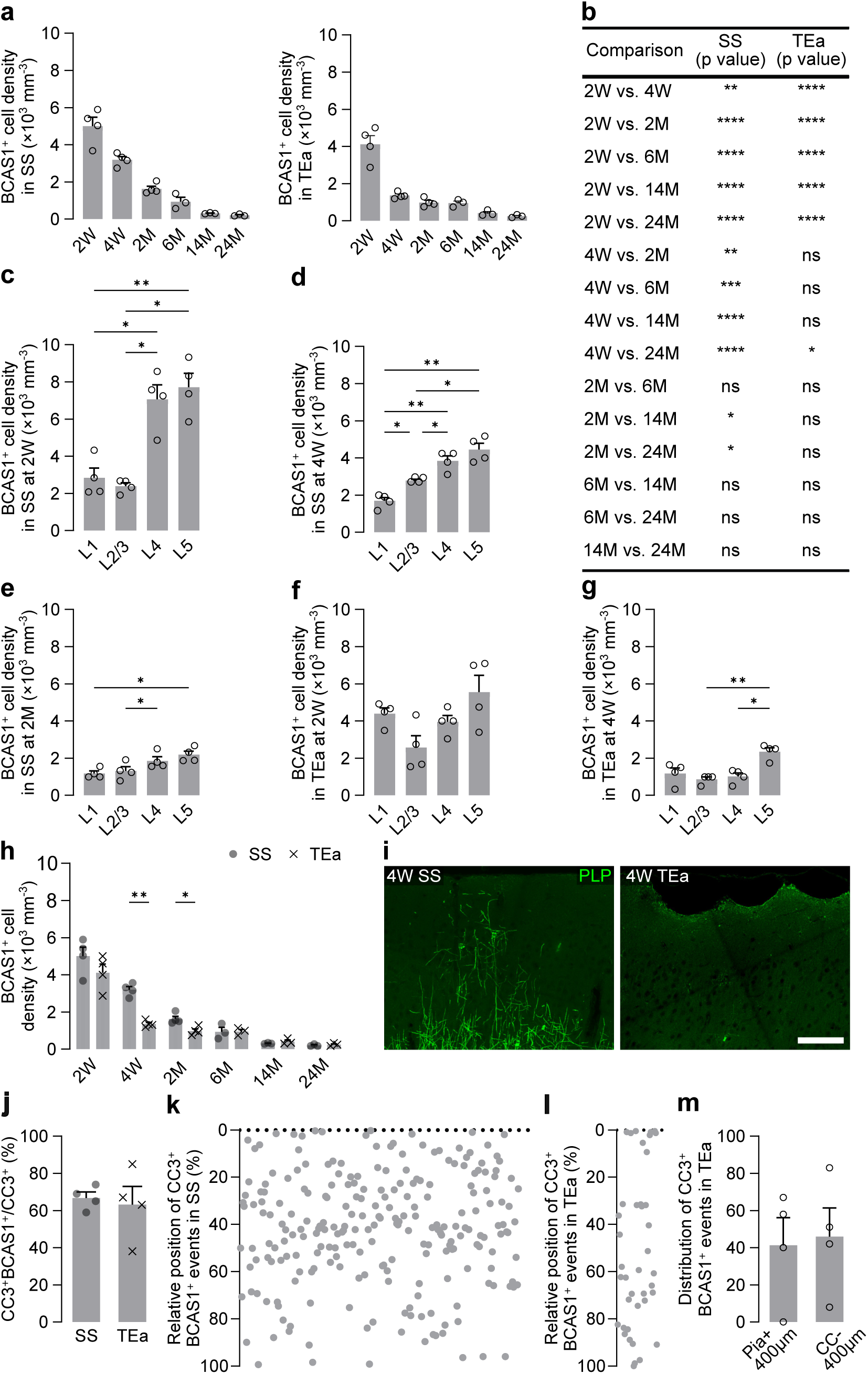
Spatiotemporal distribution of BCAS1^+^ cells and associated apoptosis. a, BCAS1^+^ cell density averaged by cortical layer in SS (left) and TEa (right) across development. b, Comparisons of BCAS1^+^ cell density averaged by cortical layer between different timepoints in SS and TEa. (a-b, 2W to 2M: n = 4 animals, 6M to 24M: n = 3 animals, one-way ANOVA with Tukey post hoc test). c-e, Cross-layer comparisons of BCAS1^+^ cell density in SS at 2W (c), 4W (d) and 2M (e) (n = 4 animals, RM ANOVA with Tukey post hoc test). f-g, Cross-layer comparisons of BCAS1^+^ cell density in TEa at 2W (f) and 4W (g) (n = 4 animals, RM ANOVA with Tukey post hoc test). h, BCAS1^+^ cell density averaged by cortical layer in SS and TEa across development (2W to 2M: n = 4 animals, 6M to 24M: n = 3 animals, multiple paired t-tests). i, Representative images for myelination in the upper 400 µm part of cortex in SS and TEa at 4W. Scale bar, 100 µm; z-projection, 20 µm. j, Percentage of CC3^+^BCAS1^+^ events among all CC3^+^ events in SS and TEa (n = 4 animals, paired t-test). k-l, Distribution of CC3^+^BCAS1^+^ events in SS (k) and TEa (l). The y-axis represents the proportional position across the full cortical depth (from pia to corpus callosum). m, Percentage of CC3^+^BCAS1^+^ events located in the upper (400 µm below the pia) and lower (400 µm above the corpus callosum) part of TEa at 2W (n = 4 animals, paired t-test). Data are presented as mean ± SEM. *P ≤ 0.05, **P ≤ 0.01.

**Supplementary figure 5.**
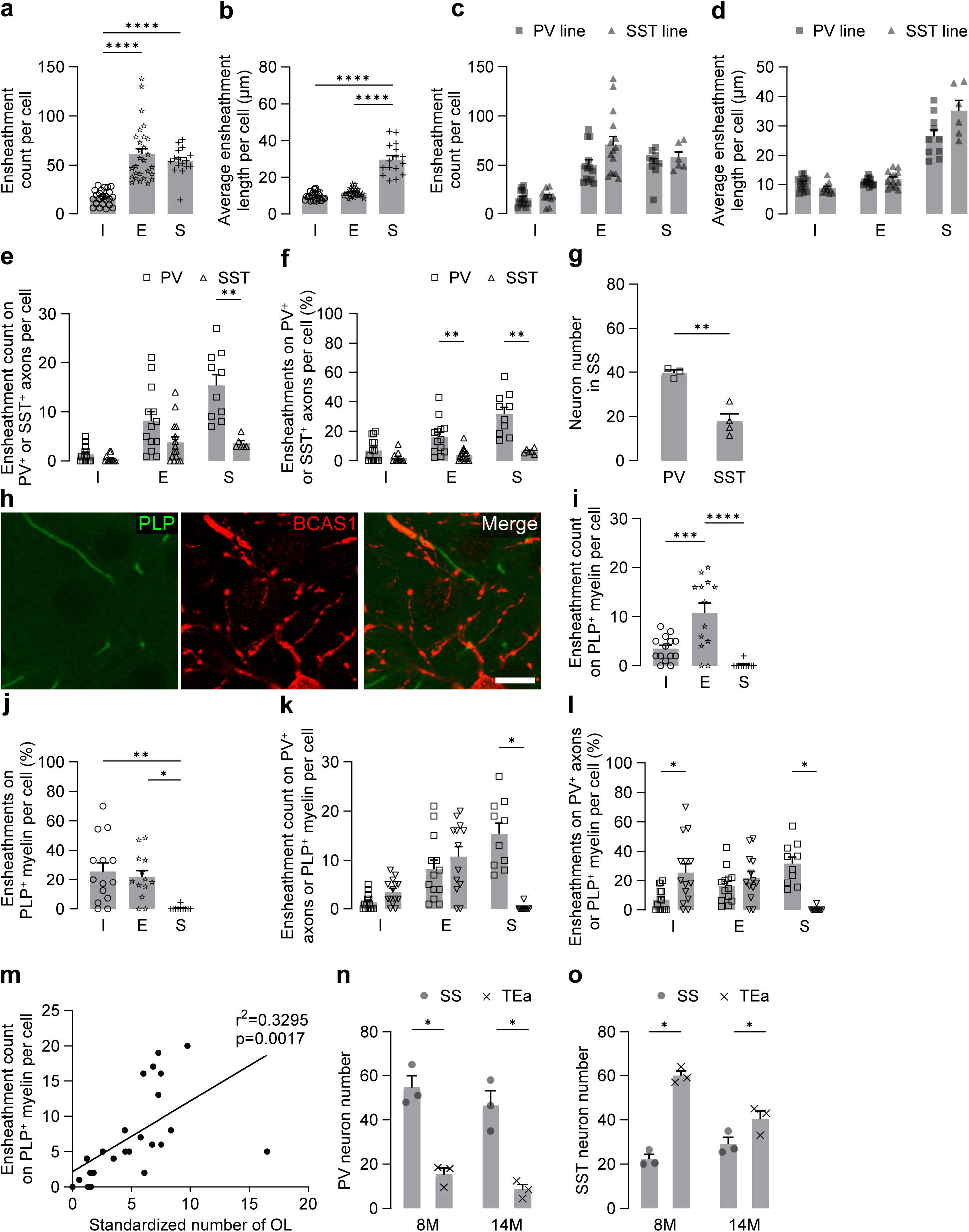
Comparison of BCAS1^+^ ensheathments on different targets. a, Number of BCAS1^+^ ensheathments for individual BCAS1^+^ cells. b, Average length of BCAS1^+^ ensheathments for individual BCAS1^+^ cells. (a-b, *initial*: n = 25 cells from 6 animals, *expansion*: n = 28 cells from 6 animals, *stabilization*: 16 cells from 6 animals, one-way ANOVA with Tukey post hoc test). c, Number of BCAS1^+^ ensheathments for individual BCAS1^+^ cells in *PV-Cre* × *tdTomato* × *Plp-GFP* mice and *SST-Cre* × *tdTomato* × *Plp-GFP* mice. d, Average length of BCAS1^+^ ensheathments for individual BCAS1^+^ cells in *PV-Cre* × *tdTomato* × *Plp-GFP* and *SST-Cre × tdTomato × Plp-GFP* mice. e, Number of BCAS1^+^ ensheathments on PV^+^ or SST^+^ axons for individual BCAS1^+^ cells. f, Percentage of BCAS1^+^ ensheathments on PV^+^ or SST^+^ axons among all the BCAS1^+^ ensheathments for individual BCAS1^+^ cells. (c-f, PV line or PV^+^ axons, *initial*: n = 14 cells from 3 animals, *expansion*: n = 13 cells from 3 animals, *stabilization*: n = 10 cells from 3 animals, SST line or SST^+^ axons, *initial*: n = 11 cells from 3 animals; *expansion*: n = 15 cells from 3 animals; *stabilization*: n = 6 cells from 3 animals, multiple t-tests). g, Number of PV and SST neurons within a 400 × 300 × 50 µm^3^ volume in layer 2/3 of SS (PV: n = 3 animals, SST: n = 4 animals, student t-test). h, Representative images of BCAS1^+^ ensheathments on PLP^+^ mature myelin. Scale bar, 10 µm; z projection, 1 µm. i, Number of BCAS1^+^ ensheathments on PLP^+^ mature myelin for individual BCAS1^+^ cells. j, Percentage of BCAS1^+^ ensheathments on PLP^+^ mature myelin among all BCAS1^+^ ensheathments for individual BCAS1^+^ cells. k, Number of BCAS1^+^ ensheathments on PV^+^ axons or PLP^+^ mature myelin for individual BCAS1^+^ cells. l, Percentage of BCAS1^+^ ensheathments on PV^+^ axons or PLP^+^ mature myelin for individual BCAS1^+^ cells. (i-l, *initial*: n = 14 cells from 3 animals, *expansion*: n = 13 cells from 3 animals, *stabilization*: n = 10 cells from 3 animals, multiple paired t-tests) m, Linear regression between the number of BCAS1^+^ ensheathments on PLP^+^ mature myelin for individual BCAS1^+^ cells and the standardized local oligodendrocyte number. n-o, Number of PV (n) and SST neurons (o) within a 400 × 300 × 50 µm^3^ volume in SS and TEa at 8M and 14M (n = 3 animals, paired t-test). Data are presented as mean ± SEM. \**P* ≤ 0.05, \*\**P* ≤ 0.01, \*\*\*\**P* ≤ 0.0001.

